# Insm1a regulates motor neuron development in zebrafish

**DOI:** 10.1101/154518

**Authors:** Jie Gong, Xin Wang, Chenwen Zhu, Xiaohua Dong, Qinxin Zhang, Xiaoning Wang, Xuchu Duan, Fuping Qian, Yu Gao, Qingshun Zhao, Renjie Chai, Dong Liu

**Author notes:** these authors contributed equally to this work (Jie Gong, Xin Wang). authors for correspondence; Contact details for Correspondence: Dong Liu, Ph.D Jiangsu Key Laboratory of Neuroregeneration, Nantong University Qixiu Road 19, Nantong, China, 226001.

## Abstract

Insulinoma-associated1a (Insm1a) is a zinc-finger transcription factor playing a series of functions in cell formation and differentiation of vertebrate central and peripheral nervous systems and neuroendocrine system. However, its roles on the development of motor neuron have still remained uncovered. Here, we provided evidences that *insm1a* was a vital regulator of motor neuron development and provide a mechanistic understanding of how it contributes to this process. Firstly, we showed the localization of *insm1a* in spinal cord and primary motor neurons (PMNs) of zebrafish embryos by *in situ* hybridization and imaging analysis of transgenic reporter line *Tg(insm1a: mCherry) ^ntu805^*. Then we demonstrated that the deficiency of *insm1a* in zebrafish larvae lead to the defects of PMNs development, including the reduction of caudal primary motor neurons (CaP) and middle primary motor neurons (MiP), the excessive branching of motor axons, and the disorganized distance between adjacent CaPs. Additionally, knockout of *insm1* impaired motor neuron differentiation in the spinal cord. Locomotion analysis showed that *insm1a*-null zebrafish significantly reduced the swimming activity. Furthermore, we proved that the *insm1a* loss of function significantly decreased the transcripts levels of both *olig2* and *nkx6.1*. Microinjection of *olig2* and *nkx6.1* mRNA rescued the motor neuron defects in *insm1a* deficient embryos. Taken together, these data indicate that *insm1a* regulates the motor neuron development, at least in part, through modulation of the expressions of *olig2* and *nkx6.1*.

## Introduction

In vertebrates, motor neurons have precise subtype identities that characterized by a number of morphological criteria, such as soma location and shape, axon path and target muscle innervation (Lewis and Eisen, 2003; Shirasaki and Pfaff, 2002). Meanwhile, motor neurons generally extend their axonal trajectory with a highly stereotyped manner during the nervous system development (Eisen, 1991; Palaisa and Granato, 2007). It has been reported that in chick and bullfrog, their motor neurons axons always followed the conservative pathways in order to project to appropriate regions of target musculatures (Farel and Bemelmans, 1985; Landmesser, 1980). In the embryo and larva of zebrafish, there are two different kinds of spinal motor neurons, which are called primary motor neurons (PMNs) and secondary motor neuron (SMNs) (Myers, 1985; Myers et al., 1986). The PMNs can be further classified into three groups, caudal primary motor neurons (CaP), middle primary motor neurons (MiP) and rostral primary motor neurons (RoP), by the positions of somata in the spinal cord and the trajectory of neuron axons (Myers et al., 1986; Westerfield et al., 1986). CaPs, whose somata locate in the middle of each spinal cord hemisegment, can innervate ventral axial muscle, and have been well studied because of their easy observation and distinct axon projection (Myers et al., 1986; Rodino-Klapac and Beattie, 2004). MiPs project axons to innervate the dorsal axial muscle, while RoPs project axons to control the middle muscle (Rodino-Klapac and Beattie, 2004). Although the somata of the three identifiable PMNs localized in different position in the spinal cord, their axons pioneer to the myoseptum through a mutual exit point (Eisen et al., 1986). Due to the identifiability of the three kinds of PMNs, they have already become an excellent system to study motor axon guidance and their intraspinal navigation (Beattie et al., 2002).

The insulinoma-associated 1 (*insm1*) gene, which is first isolated from an subtraction cDNA library of insulinoma tumor cells, encodes a DNA-binding zinc finger transcription factor with SNAG repressor motifs in N-terminal as well as Cys2-His2 Zn finger motifs in C-terminal and widely expresses in the developing nervous system, endocrine cells, pancreatic cells and related neuroendocrine tumor cells (Goto et al., 1992; Jacob et al., 2009; Jia et al., 2015b; Lan and Breslin, 2009; Xie et al., 2002). Consequently, extensive studies focused on the biological function of Insm1 in nervous and endocrine cell proliferation, differentiation and transformation have been reported in the model organisms (Farkas et al., 2008; Jia et al., 2015a; Jia et al., 2015b; Lan and Breslin, 2009; Ramachandran et al., 2012; Wildner et al., 2008). For example, in the *INSM1* knockout mice, its endocrine progenitor in the developing pancreas were less differentiated, meanwhile hormone production and cell migration were also exhibited seriously defects (Osipovich et al., 2014). Farkas et al. reported that compared to the wild type and heterozygous mice, the number of basal progenitors in the *INSM1* null dorsal telencephalon (dTel) was decreased almost half, and the radial thickness of dTel cortical plate as well as the Neurogenesis in the neocortex were also predominantly reduced after lacking *INSM1* gene (Farkas et al., 2008). In the zebrafish, *insm1a* can regulate a series of related genes, which are necessary for the Müller glia (MG) formation and differentiation as well as the zone definition of injury-responsive MG to participate in the retina regeneration (Ramachandran et al., 2012). Moreover, it was also reported that during the development of zebrafish retina, *insm1* could regulate cell cycle kinetics and differentiation of the progenitor cells by acting the upstream of the basic helix-loop-helix (bHLH) transcription factors and the photoreceptor specification genes (Forbes-Osborne et al., 2013). Although insm1a is widely detected in the nervous system and its necessity in the brain and retina development have been also illuminated well, little is known about the function and molecular mechanisms of *insm1a* in the formation and development of other neuronal types, especially in the zebrafish.

The zebrafish has become an excellent model system to investigate the mechanisms of the neuron formation and its axonal path finding due to the accessible observation of motor neurons from the initial stages of embryo development (Zelenchuk and Bruses, 2011). Here, we examined the function of *insm1a* in the primary motor neurons development by CRISPR/ Cas9-mediated knockout in the *Tg(mnx1: GFP)^ml2^* transgenic zebrafish lines and investigated the possible transcriptional network during this process.

## Materials and methods

### Zebrafish line and breeding

The zebrafish embryos and adults were maintained in zebrafish Center of Nantong University under conditions in accordance with our previous protocols (Wang et al., 2016; Xu et al., 2014). The transgenic zebrafish line, *Tg(mnx1: GFP)^ml2^*, have been described in the previous work (Zelenchuk and Bruses, 2011).

### RNA isolation, reverse transcription and quantitative PCR

Total RNA was extracted from zebrafish embryos by TRIzol reagent according to the manufacturer’s instructions (Invitrogen, USA). Genomic contaminations were removed by DNaseI, and then 2 μg total RNA was reversely transcribed using a reversed first strand cDNA synthesis kit (Fermentas, USA) and stored at -20 °C. qRT-PCR was performed using the corresponding primers (Supplementary table 1) in a 20 μl reaction volume with 10 μl SYBR premix (Takara, Japan) and *elongation factor 1a* (*ef1a*) was used as the internal control. All samples were analyzed in triplicate.

### Whole mount *in situ* hybridization

A 501 bp cDNA fragment of *insmla* was amplified from the cDNA library that established from wild type (WT) AB embryos using the specific primers of *insmla* F1 and *insmla* R1 (Supplementary table 1). Digoxigenin-labeled sense and antisense probes were synthesized using linearized pGEM-T-easy vector subcloned with this *insm1a* fragment by *in vitro* transcription with DIG-RNA labeling Kit (Roche, Switzerland). Zebrafish embryos and larvae were collected and fixed with 4% paraformaldehyde (PFA) in phosphate-buffered saline (PBS) for one night. The fixed samples were then dehydrated through a series of increasing concentrations of methanol and stored at -20°C in 100% methanol eventually. Whole mount *in situ* hybridization was subsequently performed as described in the previous study (Huang et al., 2013).

### Establishment of *Tg(insm1a: EGFP)* and *Tg(insm1a: mCherry)* transgenic line

Transgenic zebrafish were created using the Tol2kit transgenesis system and Gateway vectors. The *insm1a* promoter was cloned and insert into the p5E-MCS entry vector. A multiSite Gateway vector construction reaction (Invitrogen, USA) was conducted with the resulting p5E-insm1a together with pME-EGFP (or mCherry) and p3E-polyA subcloned into the pDestTol2pA2 to produce *insmla: EGFP* or *insmla: mCherry* construct. Subsequently, this construct was co-injected with *tol2-transposase* mRNAs into zebrafish one to two-cell-stage embryos to create the *Tg(insm1a: EGFP)^ntu804^ and Tg(insm1a: mCherry) ^ntu805^* transgenic line.

### sgRNA/ Cas9 mRNA synthesis and injection

Cas9 mRNA was obtained by *in vitro* transcription with the linearized plasmid pXT7-Cas9 following the previous methods, while sgRNAs were transcribed from the plasmid pT7 subcloned with templates that amplified by PCR with the specific forward primers and a universal reverse gRNA primer (Supplementary table 1) (Chang et al., 2013; Qi et al., 2016). Transgenic zebrafish lines *Tg(mnx1: GFP)^ml2^*, were natural mated to obtain embryos for microinjection. One to two-cell stage zebrafish embryos was injected with 2-3 nl of a solution containing 250 ng/μl Cas9 mRNA and 15 ng/μl sgRNA. At 24 hours post fertilization (hpf), zebrafish embryos were randomly sampled for genomic DNA extraction according to the previous methods to determine the indel mutations by sequencing.

### Morpholino and mRNAs injections

Translation blocking antisense Morpholino (MOs; Gene Tools) against the ATG-containing sequence was designed (5’-AAATCCTCTGGGCATCTTCGCCAGC-3’) to target the translation start site according to the manufacturer’s instruction and the other MO oligo (5’-CCTCTTACCTCAGTTACAATTTATA-3’) was used as standard control. The MOs were diluted to 0.3mM with RNase-free water and injected into the yolk of one to two-cell stage embryos and then raised in E3 medium at 28.5 °C.

The cDNAs containing the open reading frame of the two genes were cloned into PCS2^+^ vector respectively and then were transcribed *in vitro* using the mMESSAGE mMACHIN Kit (Ambion, USA) after the recombinant plasmids linearized with NotI Restriction Enzyme (NEB, England), and then the capped mRNAs were purified by RNeasy Mini Kit (Qiagen, Germany). 2nl target genes and *mCherry* mRNA mixture (1:1) was injected at 20ng/μl into 1/2-cell stage embryos.

### Locomotion analysis in zebrafish larvae

To determine whether the deficiency of *insmla* affect spontaneous movement, knockout and normal larvae respective were raised in a 24-well culture plate with one larva in each well filled with 1 ml E3 medium. The 24-well culture plate was transferred to the Zebralab Video-Track system (Zebrabox, France) equipped with a sealed opaque plastic box that kept insulated from laboratory environment, an infrared filter and a monochrome camera. After adapting for 30 min, travelled distances of the larvae were videotaped with every 5 mins forming a movement distance and trajectory by the linked software.

### Microscopy and statistical analysis

Zebrafish embryos were anesthetized with E3/0.16 mg/mL tricaine/1% 1-phenyl-2-thiourea (Sigma, USA) and embedded in 0.8% low melt agarose, and then were examined with a Leica TCS-SP5 LSM confocal imaging system. For the results of *in situ* hybridization, Photographs were taken using an Olympus stereomicroscope MVX10. Statistical comparisons of the data were carried out by student’s t-test or one-way analysis of variance (ANOVA) followed by Duncan’s test, and P values < 0.05 were considered statistically significant. All statistical analysis was performed using the SPSS 13.0 software (SPSS, USA).

## Result

### *Insm1a* is expressed in spinal cord and PMNs of Zebrafish

To analyze the expression of *insm1a* in zebrafish nervous system, we performed the whole amount *in situ* hybridization (WISH) analysis with a digoxigenin-labeled *insmla* probe. Similar to the previous study (Lukowski et al., 2006), at late somitogenesis (24 hpf) *insm1a* transcripts were apparently localized in ventral part of the neurons in the spinal cord, where most of the motor neurons locate at this stage (Figure 1A).

**Figure 1.**
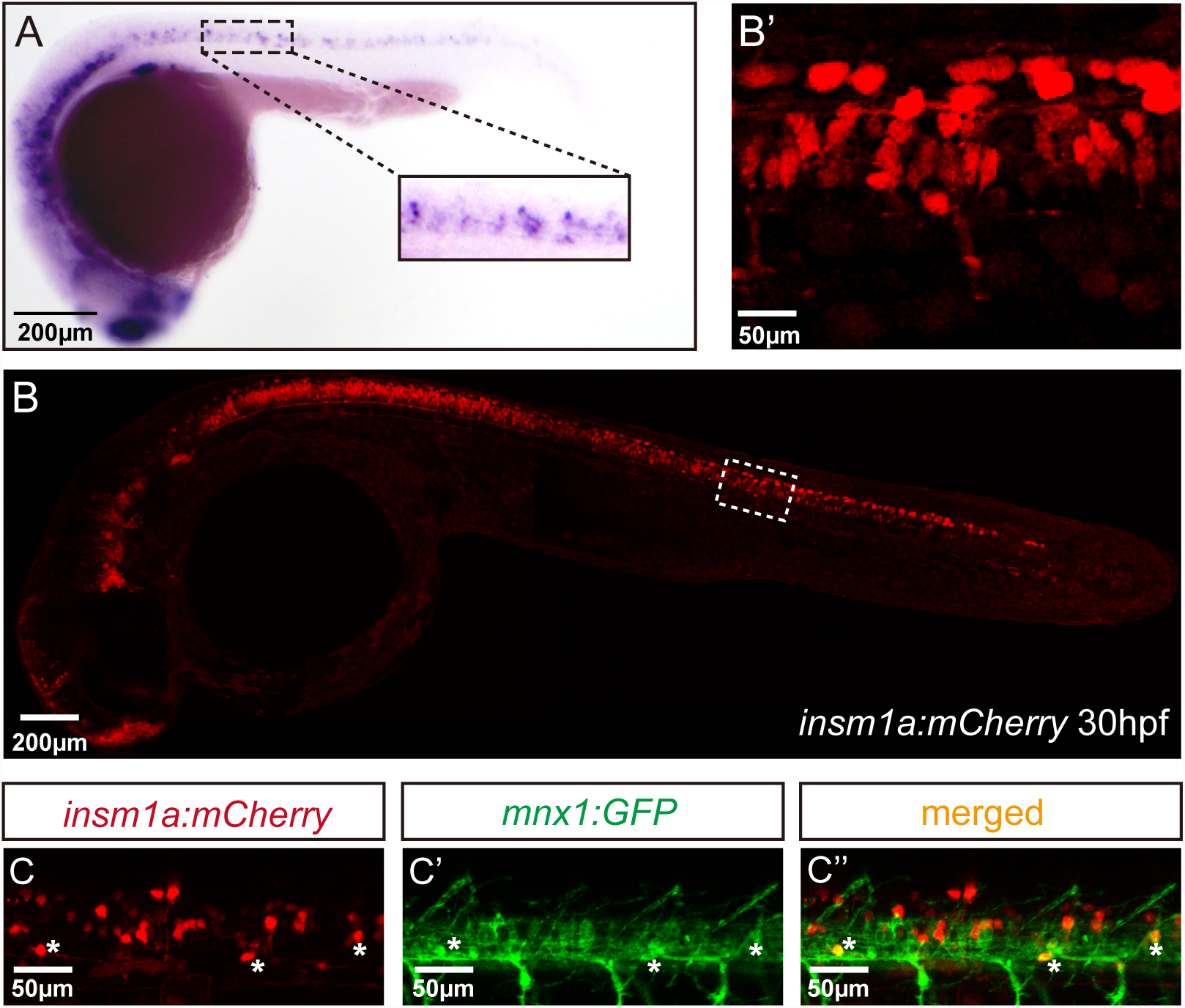
*Insm1a* expression in embryonic zebrafish spinal cord and primary motor neurons. A. At 24 hpf, the *in situ* hybridization signal of insm1a is localized in the spinal Scale bar = 200 μm. The confocal imaging analysis of the transgene *insm1a: mCherry* expression at 30 hpf. Square in dash line indicates the magnified region in B’. Scale bar = 50 m. C, C’ and C”. Confocal imaging analysis of *Tg(mnx1: GFP)^ml2^ Tg(insm1a: mCherry) ^ntu805^* transgenic line.

To further determine the localization of *insm1a*, we generated the *Tg(insm1a: EGFP) ^ntu804^* and *Tg(insm1a: mCherry) ^ntu805^* transgenic zebrafish lines, in which the *insm1a* promoter driving expression of EGFP or mCherry respectively. It was shown that at 30 hpf the *insm1a: mCherry* and *insm1a:EGFP* expression was observed in the spinal cord, retina and brain, that was similar with the results of *in situ* hybridization (Figure 1B, B’; Supplementary Figure 1A, A’) (Lukowski et al., 2006). In addition, we found that *insm1a:EGFP* expression was highly activated in Mu□ller glia of injury sites in adult zebrafish retina (Supplementary Figure 1B), which was consistent with the ISH data carried out by Rajesh Ramachandran et al (Ramachandran et al., 2012). These results suggest that the transgenes recapitulate the endogenous *insm1a* expression.

To investigate whether *insm1a* is expressed in motor neurons, we outcrossed *Tg(insm1a: mCherry) ^ntu805^* transgenic line with *Tg(mnx1: GFP)^ml2^* line, in which GFP was motor neurons labeled by under the control of *mnx1* promoter (Zelenchuk and Bruses, 2011). We found that the GFP+ motor neurons are also labeled with mCherry fluorescence (Figure 1C-C”), suggesting *insm1a* is expressed in motor neurons.

### Knockout of *insm1a* caused primary motor neurons developmental defect

In order to examine whether *insm1a* was required for the development of motor neuron, the CRISPR/Cas9 system was utilized to knockout *insm1a* in *Tg(mnx1: GFP)^ml2^* transgenic zebrafish line. To ensure complete disruption of functional proteins, we chose the target sites near the translation start codon (ATG) in the exon1 of zebrafish *insm1a* (Supplement Figure 2A). The selected gRNA-Cas9 system efficiently induced mutations in the targeting site with 4 types of mutations were identified (Supplementary Figure 2B). The mutated alleles included a 5-bp deletion, an 8-bp deletion and two 10-bp deletion, which all result in reading frame shift and premature translation termination (Supplementary Figure 2C). And the 8-bp deletion mutant line was used for the following experiments.

It was observed that *insm1a* knockout caused significant developmental defect of motor neurons(Figure 2). Firstly, the number of MiPs and CaPs were significantly reduced in the *insm1a* mutants (Figure 2A). We counted the number of Caps and classified the zebrafish embryo into 3 categories by its defective degree: severe group with over 80% loss of Caps, moderate group with less than 80% loss and normal group with less than 20% loss (In the following statistical analysis, the zebrafish with less than 20% loss was defined as normal, whereas, it was abnormal). These results revealed that 48.1% severe and 32.1% moderate defect were found in the *insm1a* mutants, while there was only 7.9% moderate defect in the control group (Figure 2B). Similarly, the MiPs were also obviously impaired in *insmla* knockout embryos (Fig. 2A, F). Moreover, we found that these abnormal phenotypes of motor neurons could not recover at later stages we checked (Supplementary Figure 3).

**Figure 2.**
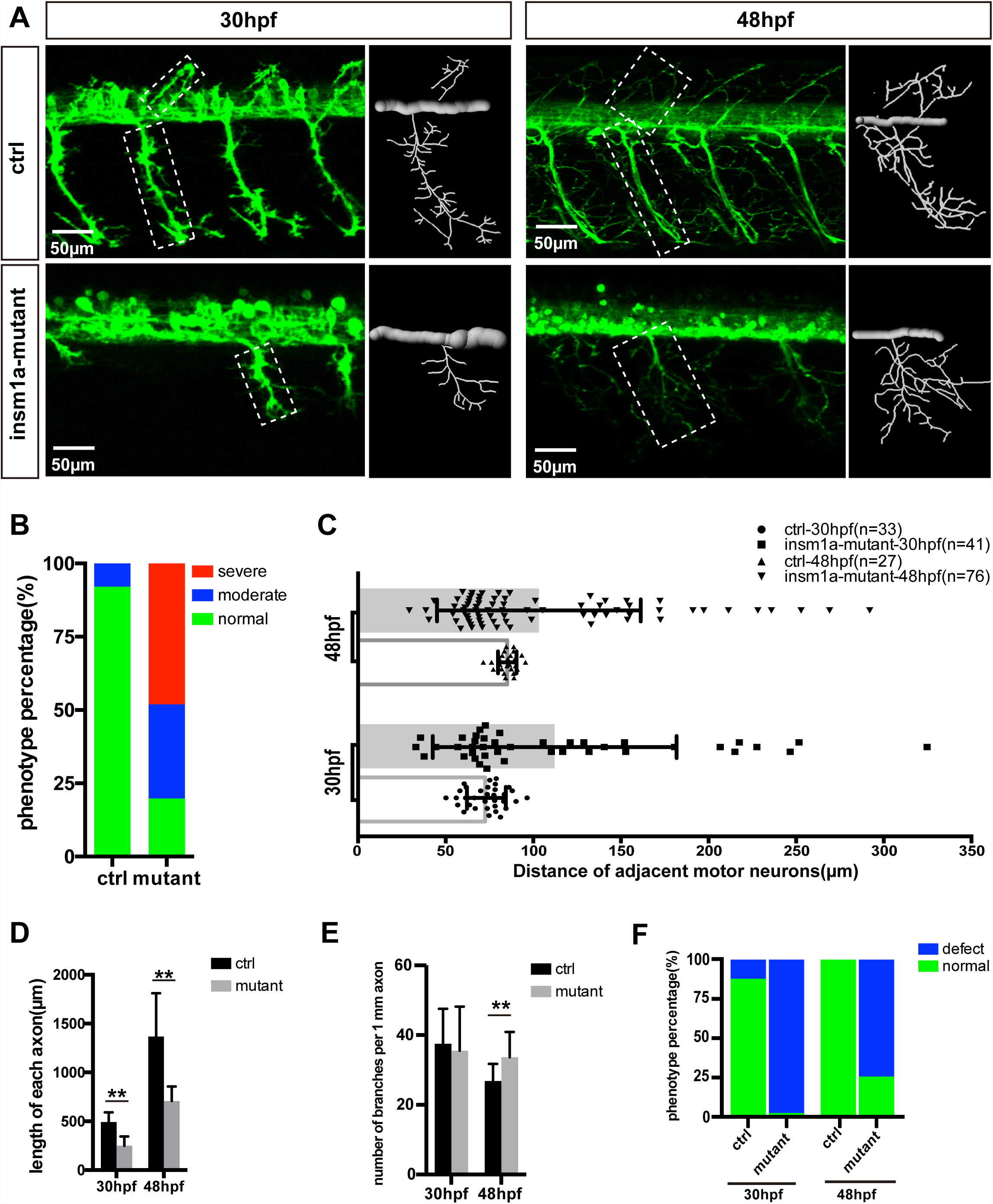
Primary motor neuron morphogenesis defects in the *insm1a* mutant zebrafish embryos. A. Confocal imaging analysis of primary motor neuron in control group and *insmla* mutant groups at 30 hpf and 48 hpf *Tg(mnx1: GFP)^ml2^*. Caps in dash line are showed in diagrams. Scale bar= 50 μm. B. Quantification of zebrafish embryos with abnormal Caps. The zebrafish embryos are classified into three categories according to its loss degree: severe group with over 80% loss of Cap primary motor neuron, moderate group with less than 80% loss and normal group with less than 20% loss. C. Quantification of distance between adjacent motor neurons (μm) in control group and *insm1a* mutant groups at 30 hpf (n=33 and 41 respectively) and 48 hpf (n=27 and 76 respectively). D and E. The length and branching number of Cap axons in control group and *insmla* mutant groups at 30 hpf and 48 hpf. Asterisks above the bars are significantly different (*P*<0.05). F. Quantification of zebrafish embryos with abnormal Caps at 30 hpf and 48 hpf.

Moreover, the morphology of motor neurons was significantly affected in the *insm1a* mutants (Figure 2A, D and E). The axons of Caps in *insm1a* mutants were shorter and failed to reach the ventral musculatures. The branches density of the Caps in *insm1a* mutants is higher than that of the Caps in control. In the *insm1a* mutants, there are around 34 branch points of per 1 mm CaP axon at 48 hpf, only 31 in control embryos (Figure 2E). With the larvae development, the excess branching became more and more pronounced (Supplement Figure 3). In addition, statistical analysis revealed that the average length of each CaP anon in the *insm1a* mutants was 707.9 um at 48 hpf, while in control embryos it increased to 1367.9 um (Figure 2D). Interestingly, we also found that the distances between adjacent CaPs were significantly variant in *insm1a* mutants (Figure 2C).

In order to validate the developmental defects of motor neuron was specifically caused by the *insm1a* inactivation, further experiments were carried out. The embryos that injected with an *insm1a* translation blocking morpholino displayed the similar motor neuron with that observed in the *insm1a* mutants (Supplement Fig. 4). To confirm phenotypic specificity induced by the *insm1a* MO injection, we performed rescue experiment by co-injection of 50 ng of *insm1a* mRNA with *insm1a* MO into zebrafish embryos, and the results showed that the co-injection significantly decreased the loss and premature branching of PMNs (Fig. 4E). Taken together, these results indicate those motor neuron developmental defects were caused by loss of *insm1a*.

**Figure 4.**
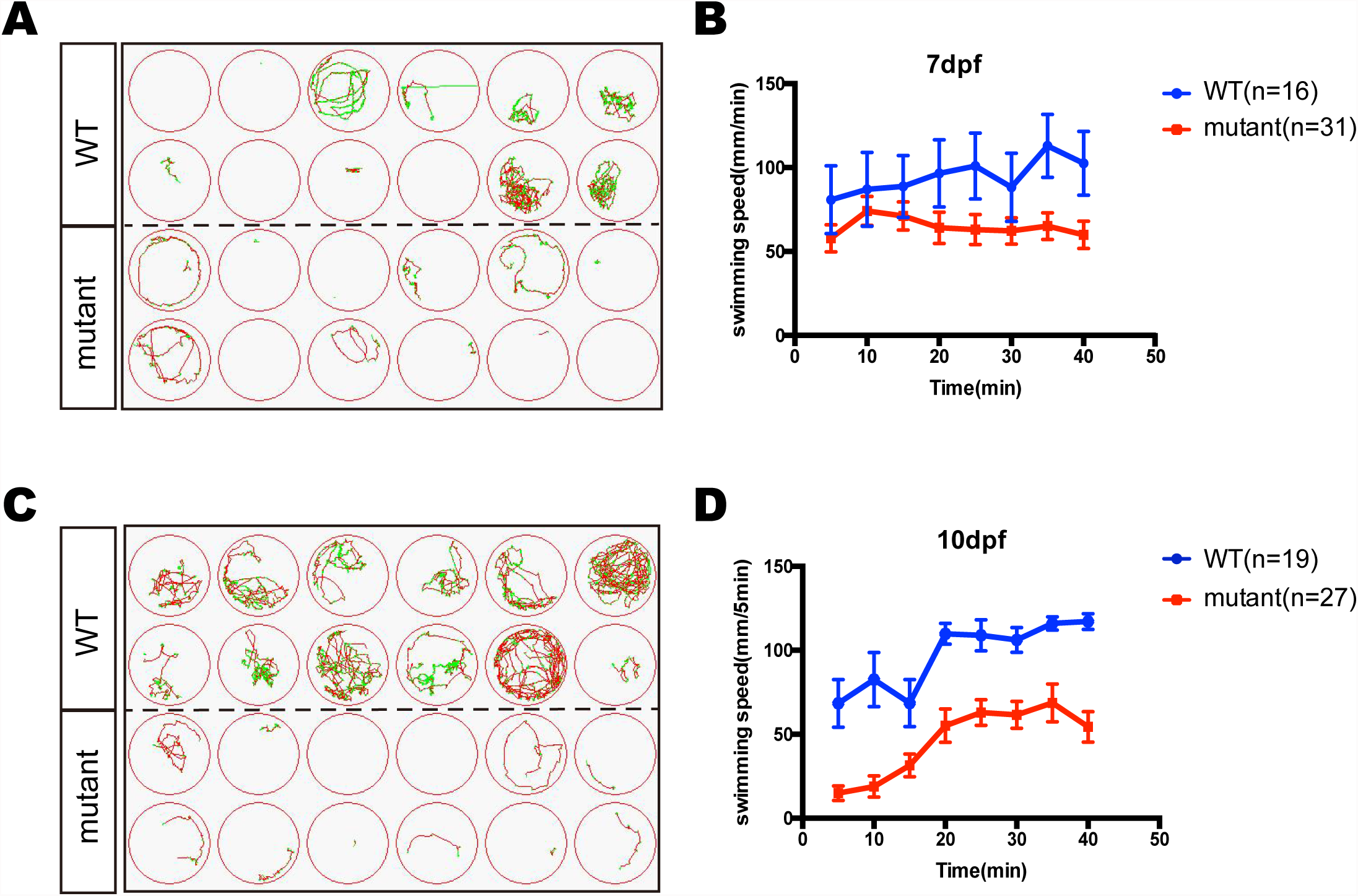
The swimming behavior analysis of control and *insm1a* mutant zebrafish embryos at 7 dpf and 10 dpf. A and C. The swimming trajectory of the control and *insm1a* mutant zebrafish embryos at 7 dpf and 10 dpf. B and D. Quantification of the swimming distance of control and *insm1a* mutant zebrafish embryos at 7 dpf and 10 dpf per 5 mins (n = 36 in each group). Asterisks indicate the statistically significant difference compared with the control (P < 0.05).

### Insm1a deficiency suppressed neuronal cells differentiation

The confocal imaging analysis discovered that there are a number of round and not well differentiated GFP positive cells in *Tg(mnx1: GFP)^ml2^ insm1a* mutants (Figure 3A). Statistical analysis showed that at 30 and 48 hpf the number of these undifferentiated cell in the *insm1a* deficiency zebrafish was significantly higher than that in the control fish (Fig. 3B). We also observed these undifferentiated cells in *insm1a* morphants, however the number is less than that in mutants (Figure 3A, B).

**Figure 3.**
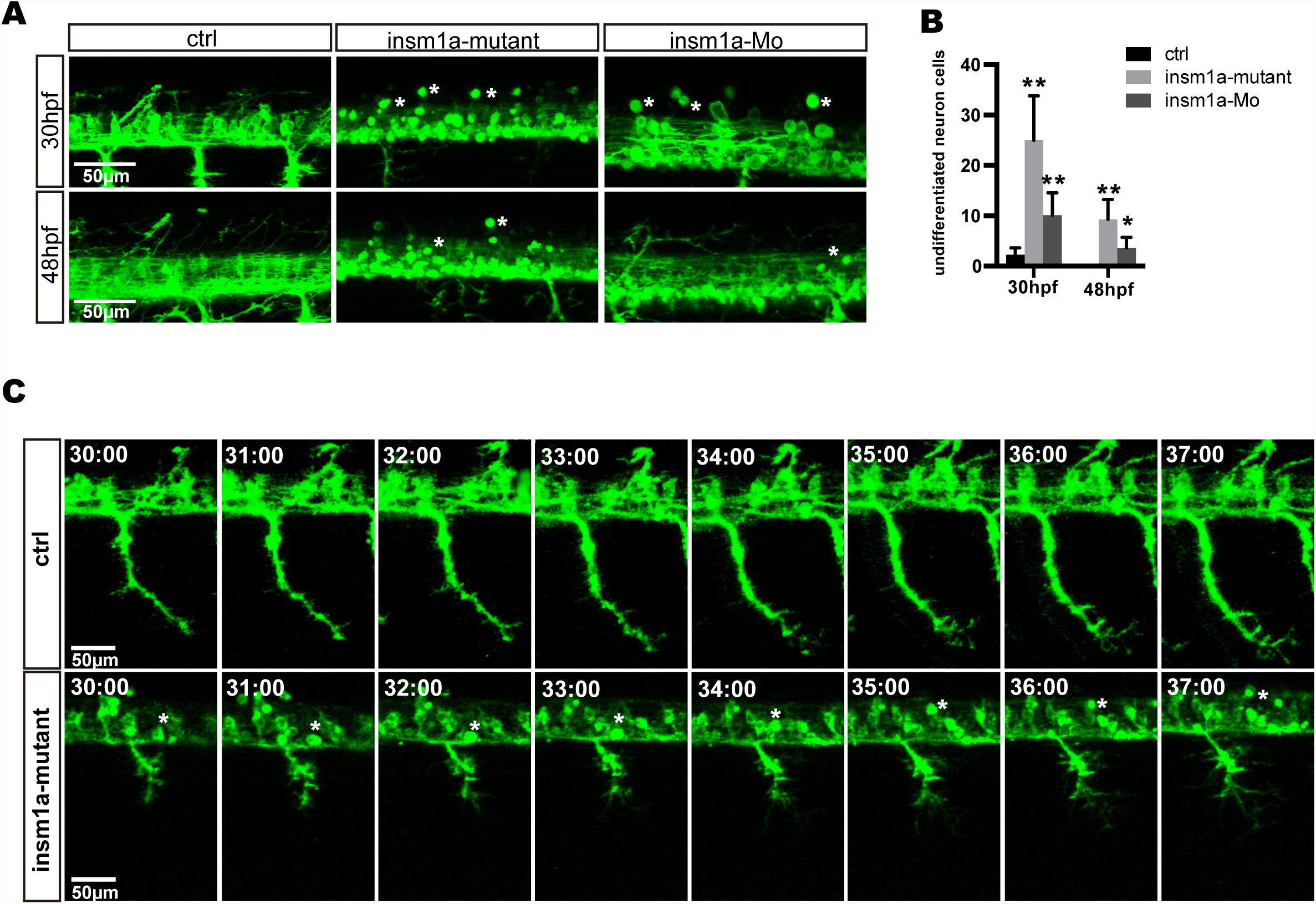
*Insm1a* deficiency suppressed neuronal cells differentiation. A. Confocal imaging analysis of primary motor neuron in control group, *insm1a* mutant group and morphant group at 30 hpf and 48 hpf *Tg(mnx1: GFP)^ml2^*. Phenotypes of neuronal cells in the spinal cord in control group, morphant group and insm1a mutant groups at 30 hpf and 48 hpf. Asterisks indicate undifferentiated neuronal cells. Scale bar = 50 μm. B. Quantification of the undifferentiated neuronal cell in the *insm1a* different treatment zebrafish. Asterisks above the bars are significantly different (P<0.05). C. Time-lapse imaging analysis of the primary motor neuron in control group and insm1a mutant groups. Asterisks represent undifferentiated neuronal cells. Scale bar = 50 μm

To further investigate the cellular mechanism underlying the motor neuronal defects in *insm1a* deficient embryos, we performed confocal time-lapes imaging analysis. It was found that in control embryos the axon of CaP sprouted from the spinal cord, and extended towards to the ventral muscle (Figure 3C). In control embryos the axon of CaP start to branch when it across the midline, while in *insm1a* mutants the axon initiate to branch once it come out from the spinal cord (Figure 3CA and Supplementary Movie1, 2). In addition, we found that those round GFP positive cells did not develop neuronal projections (Figure 3CA and Supplementary Movie 1).

### Knockout of *insm1a* reduced the zebrafish swimming activity

In order to investigate whether the motor neuron defects affect the motor ability, *insm1a* mutant zebrafish larvae were further performed for 40-min free-swimming activity test independent of any stimuli at 7 and 10 dpf. It was demonstrated the movement trajectory and swimming distance per 5 mins, which could reflect the swimming speed, of insm1a mutant zebrafish larvae were significantly decreased compared to that in the controls at both 7 dpf and 10 dpf (Figure 4). The movie in the supplementary material showed that swimming behavior could be easily discovered in the control group, while the zebrafish in mutant group keep involuntomotory (Supplementary movie 3). Additionally, we also discovered that under the stereoscopic microscope the mutant zebrafish became insensitive to the touch response (data not shown).

### The *insm1a* deficiency caused alteration of gene expression involved in motor neuron development

Since Insm1a is a transcription factor, we supposed that motor neuron developmental defects in *insm1a* deficient embryos were associated with altered expression of genes downstream of *insm1a* or the genes participating in the motor neuron development. Based on the previous studies, *NNR2a*, *NNR2b*, *islet2*, *Asci1a*, *Asci1b*, *shh*, *Ngn2*, *Nkx6.1* and *olig2* were selected to do the qRT-PCR analysis in wild-type (WT) and Insm1 deficiency zebrafish embryos. The results showed that expressions of *NNR2a*, *NNR2b*, *islet2*, *Asci1a* and *Asci1b* were significantly influenced in the *insm1a* deficiency zebrafish compared to the controls (Supplement Figure 5). We also found that the expression of *shh* was obviously elevated in *insm1a* mutants at 19, 24 and 36 hpf (Supplement Figure 5). Interestingly, *olig2* and *nkx6.1* transcripts dramatically decreased in *insm1a* deficient embryos.

**Figure 5.**
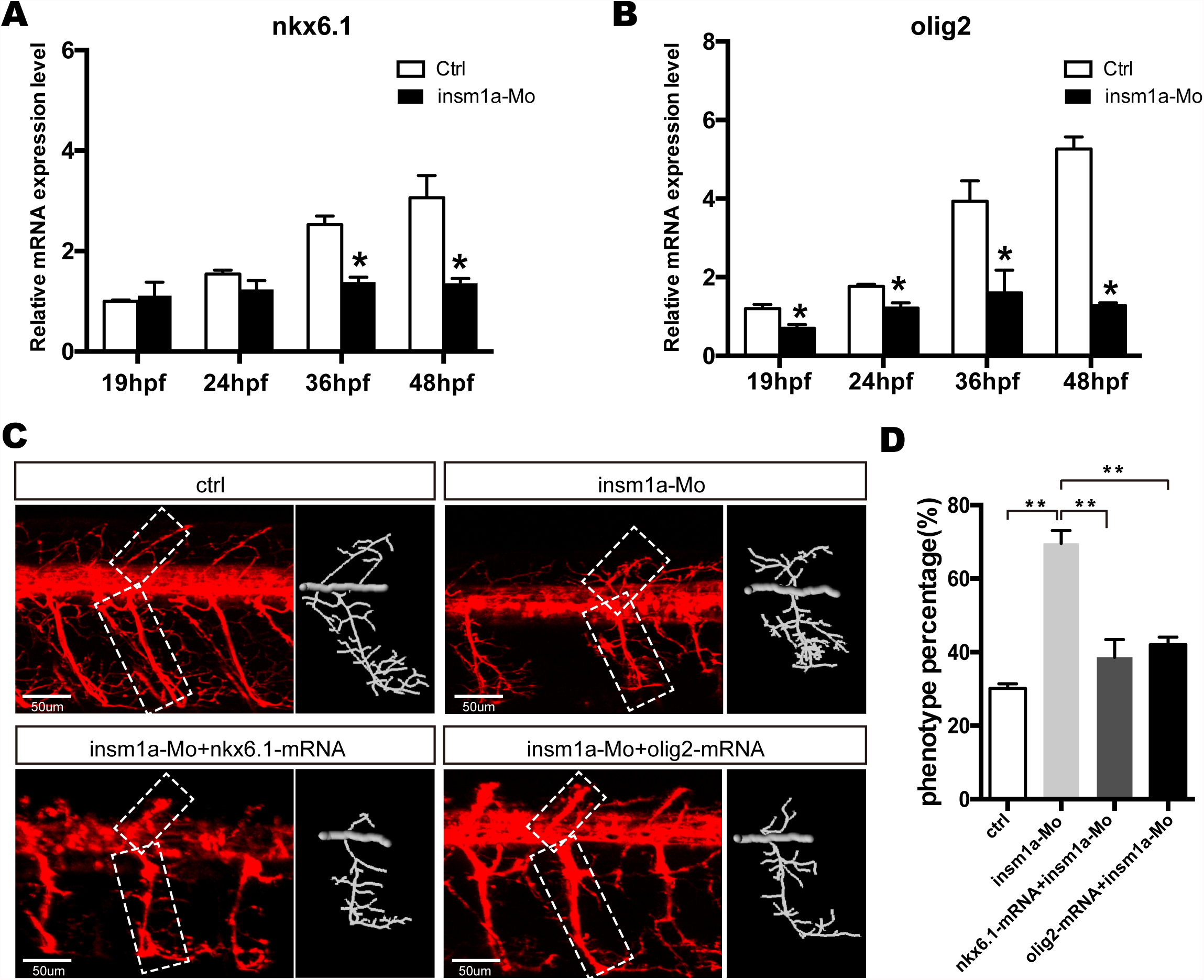
Over expressions of *nkx6.1* and *olig2* rescued the motor neuron defects in *insm1a* deficient embryos. A and B. Effects of *insm1a* knockdown on the expressions of *nkx6.1* and *olig2* at 19, 24, 36 and 48 hpf. Asterisks indicate significant differences compared with the control (P<0.05). C. Abnormal Caps in insm1a knockdown zebrafish embryos were restored by co-injection of *nkx6.1* or *olig2* mRNA. Diagrams of Caps in dash line are displayed near the corresponding confocal image. Scale bar = 50 μm. D. Quantification of zebrafish embryos with abnormal Cap primary motor neuron. Asterisks indicate significant differences compared with the control (P<0.05).

### *Olig2* and *nkx6.1* over expression rescued the motor neuron defects in *insm1a* deficient embryos

As the downergulation of *olig2* and *nkx6.1* in *insm1a* loss of function embryos, we reasoned that Insm1a might bind the transcriptional regulatory elements of these two genes. Based on the JASPAR 2016 database(Mathelier et al., 2016) analysis, we found that both *olig2* and *nkx6.1* contained the putative binding sites of Insm1a (Supplement Figure 6), suggesting Insm1a directly regulates the expression of *olig2* and *nkx6.1* during PMNs development. To investigate whether the motor neuronal defects in *insm1a* deficient embryos were caused by reduced expression of *olig2* and *nkx6.1*, we try to rescue the phenotype with olig2 and n kx6.1 gain of function in *insm1a* deficient embryos. It was shown that coinjecting both *olig2 and nkx6.1* mRNA respectively with insm1a MO significantly reduced the motor neuronal defects caused by loss of *insm1a* (Figure 5C, D). of 69.6% zebrafish embryos injected with *insm1a* MO at 48 hpf had the motor neuron developmental defects, while only 42.1% had the motor neuronal phenotype in the *olig2* mRNA and *insm1a* MO co-injection group (Figure 5D). Similarly, after *nkx6.1* mRNA and *insm1a* MO co-injection, the ratio of motor neuronal phenotype decreased to 38.6% (Figure 5D).

## Discussion

As one of the most conserved zinc-finger transcriptional factor, Insm1a plays important roles in various biological processes in vertebrates (Forbes-Osborne et al., 2013; Jacob et al., 2009; Jia et al., 2015b; Lorenzen et al., 2015; Osipovich et al., 2014; Wildner et al., 2008). Previous studies have identified its role in regulating the endocrine cells divisions of the pancreas, the neuroendocrine development, the differentiation of retina progenitors and neurogenesis of nervous system (Duggan et al., 2008; Farkas et al., 2008; Gierl et al., 2006; Jacob et al., 2009; Lan and Breslin, 2009; Ramachandran et al., 2012). Currently, our data in this study provided with new insights into the role of Insm1a in motor neuron development.

Our WISH data and previous study (Ramachandran et al., 2012) demonstrated that *insm1a* transcripts were detected in retina and spinal cord at 24 hpf. Furthermore, imaging analysis of our established transgenic reporter line *Tg(insm1a: EGFP)^ntu804^ and Tg(insm1a: mCherry) ^ntu805^* verified the results of *in situ* hybridization, and revealed the expression of GFP or mCherry that were driven by *insm1a* promoter in the PMNs. It is well known that the spinal cord contained PMNs and then project their axons out of the spinal cord to the terminal musculature with the embryo development (Davis-Dusenbery et al., 2014). Taken together, the localization data of insm1a from both *in situ* hybridization analysis and the study based on transgenic reporter line suggested that *insm1a* might participate in the regulation of PMNs development.

To test whether *insm1a* was required for formation of PMNs, we generated CRISPR/Cas9-mediated *insm1a* mutants and showed that obvious motor neuron loss and defects of the PMNs axons. Moreover, we also performed *insm1a* knockdown and the results showed similar PMNs defects as the ones produced by the *insm1a* knockout. In wild embryos, exuberant side branches developed by around 72 hpf, and then invaded into myotome to form distributed neuromuscular synapses (Downes and Granato, 2004; Liu and Westerfield, 1990). These results suggest that *insm1a* is pivotal for the primary motor axon development to block precociously extending into muscle territories. Additionally, locomotion analysis displayed a typical low activity swimming behavior in *insm1a* mutant zebrafish. It was known motor neuron is a major kind of cell type that regulated swimming behavior in zebrafish during early development (Brustein et al., 2003). Previous studies also showed a significant involvement of motor neuron in the overall locomotor behavior (Flanagan-Steet et al., 2005; Levin et al., 2009). Currently, the decrease of swimming activity in this study was consistent with the motor neuron defects in the *insm1a* knockout zebrafish.

Another prominent phenotype in the *insm1a* deficiency zebrafish was the disorganized distance between adjacent Caps, which might be caused by the ectopic departure of motor axons from the spinal cord (Palaisa and Granato, 2007). During the zebrafish PMNs development, the three kinds of PMN axons firstly longitudinally migrated towards a segmental spinal cord exit point, and then they diverged to individual-specific trajectories (Eisen et al., 1986; Myers et al., 1986). It has been reported that axonal exit sites at the spinal cord might be restricted and conserved (Niederlander and Lumsden, 1996). However, the change of distance between adjacent motor axon and the formation of abnormal axons in our study suggested that the motor axons to form exit points at any positions along entire length of spinal cord in *insm1a* mutants. The similar phenotypes were also showed in plexin A3 and semaphorin 3A morphants (Feldner et al., 2007; Palaisa and Granato, 2007; Tanaka et al., 2007). Additionally, Birely et al. also reported the phenotype that motor axons departed from the spinal cord at the ectopic points accompanied with defects in slow muscle fiber development (Birely et al., 2005). These studies suggested that the low activity swimming behavior in *insm1a* mutant zebrafish might be involved in the ectopic departure of motor axons from the spinal cord.

In this study we also found that loss function of Insm1a obviously impaired the motor neuronal differentiation. Similar role of *insm1a* was also demonstrated in the retina of zebrafish that the cells differentiation and migrating in the zebrafish injected with *insm1a* MO were significantly suppressed compared that in the control group after the retinal injury (Forbes-Osborne et al., 2013; Ramachandran et al., 2012). In vertebrates, Insm1 stimulates cell cycle exit by suppressing expression of cell proliferation related genes and relieving repression of p57kip2, a cyclin kinase inhibitor that along with p27kip1 drives cell cycle exit (Dyer and Cepko, 2001). One consequence of Insm1a driven cell cycle exit is progenitor differentiation (Ramachandran et al., 2012). The undifferentiated cells in the spinal cord of *insm1a* mutants confirm the role of this transcriptional factor in cell differentiation in more cell types.

There are a series of genes have been identified to contribute to motor neuron formation and development (Cheesman et al., 2004; Park et al., 2002). It has been reported that *nkx6.1* and *olig2* were dynamically expressed in zebrafih motor neuron and required for motor neuron development and the downregulation of the two gene lead to developmental defect of motor neuron which was similar with that in *insm1a* mutant (Cheesman et al., 2004; Hutchinson et al., 2007; Park et al., 2002). Conversely, overexpression of *nkx6.1* or *olig2* by mRNA injection could significantly promote the development of the PMNs (Hutchinson et al., 2007; Park et al., 2002). Current study revealed that the inactivation of *insm1a* resulted in the significant decrease of *nkx6.1* and *olig2* expression levels. We further showed that *olig2* and *nkx6.1* overexpression rescued the motor neuron defects in insm1a deficient embryos. These data suggested that *insmla* regulates the motor neuron development, at least in part, by regulating the expressions of *olig2* and *nkx6.1*.

## Acknowledgments

This study was supported by grants from the National Natural Science Foundation of China (31400918; 41606169; 81570447) and Natural Science Foundation of Jiangsu Province (BK20160418).

## References

Beattie, C.E., Granato, M., and Kuwada, J.Y. (2002). Cellular, genetic and molecular mechanisms of axonal guidance in the zebrafish. Results Probl Cell Differ 40, 252-269.

Birely, J., Schneider, V.A., Santana, E., Dosch, R., Wagner, D.S., Mullins, M.C., and Granato, M. (2005). Genetic screens for genes controlling motor nerve-muscle development and interactions. Dev Biol 280, 162-176.

Brustein, E., Saint-Amant, L, Buss, R.R., Chong, M., McDearmid, J.R., and Drapeau, P. (2003). Steps during the development of the zebrafish locomotor network. J Physiol Paris 97, 77-86.

Chang, N., Sun, C., Gao, L, Zhu, D., Xu, X., Zhu, X., Xiong, J.W., and Xi, J.J. (2013). Genome editing with RNA-guided Cas9 nuclease in zebrafish embryos. Cell research 23,465-472.

Cheesman, S.E., Layden, M.J., Von Ohlen, T., Doe, C.Q., and Eisen, J.S. (2004). Zebrafish and fly Nkx6 proteins have similar CNS expression patterns and regulate motoneuron formation. Development 131, 5221-5232.

Davis-Dusenbery, B.N., Williams, L.A., Klim, J.R., and Eggan, K. (2014). How to make spinal motor neurons. Development 14, 491-501.

Downes, G.B., and Granato, M. (2004). Acetylcholinesterase function is dispensable for sensory neurite growth but is critical for neuromuscular synapse stability. Dev Biol 270, 232-245.

Duggan, A., Madathany, T., de Castro, S.C., Gerrelli, D., Guddati, K., and Garcia-Anoveros, J. (2008). Transient expression of the conserved zinc finger gene INSM1 in progenitors and nascent neurons throughout embryonic and adult neurogenesis. J Comp Neurol 507,1497-1520.

Dyer, M.A., and Cepko, C.L. (2001). p27Kip1 and p57Kip2 regulate proliferation in distinct retinal progenitor cell populations. J Neurosci 21, 4259-4271.

Eisen, J.S. (1991). Determination of primary motoneuron identity in developing zebrafish embryos. Science 252, 569-572.

Eisen, J.S., Myers, P.Z., and Westerfield, M. (1986). Pathway selection by growth cones of identified motoneurones in live zebra fish embryos. Nature 320, 269-271.

Farel, P.B., and Bemelmans, S.E. (1985). Specificity of motoneuron projection patterns during development of the bullfrog tadpole (Rana catesbeiana). J Comp Neurol 238,128-134.

Farkas, L.M., Haffner, C, Giger, T., Khaitovich, P., Nowick, K., Birchmeier, C, Paabo, S., and Huttner, W.B. (2008). Insulinoma-associated 1 has a panneurogenic role and promotes the generation and expansion of basal progenitors in the developing mouse neocortex. Neuron 60, 40-55.

Feldner, J., Reimer, M.M., Schweitzer, J., Wendik, B., Meyer, D., Becker, T., and Becker, C.G. (2007). PlexinA3 restricts spinal exit points and branching of trunk motor nerves in embryonic zebrafish. J Neurosci 27, 4978-4983.

Flanagan-Steet, H., Fox, M.A., Meyer, D., and Sanes, J.R. (2005). Neuromuscular synapses can form in vivo by incorporation of initially aneural postsynaptic specializations. Development 132, 4471-4481.

Forbes-Osborne, M.A., Wilson, S.G., and Morris, A.C. (2013). Insulinoma-associated la (Insmla) is required for photoreceptor differentiation In the zebrafish retina. Dev Biol 380,157-171.

Gierl, M.S., Karoulias, N., Wende, H., Strehle, M., and Birchmeier, C. (2006). The zinc-finger factor Insm1 (IA-1) is essential for the development of pancreatic beta cells and intestinal endocrine cells. Genes Dev 20, 2465-2478.

Goto, Y., De Silva, M.G., Toscani, A., Prabhakar, B.S., Notkins, A.L., and Lan, M.S. (1992). A novel human insulinoma-associated cDNA, IA-1, encodes a protein with “zinc-finger” DNA-blnding motifs. J Biol Chem 267,15252-15257.

Huang, Y., Wang, X., Wang, X., Xu, M., Liu, M., and Liu, D. (2013). Nonmuscle myosin II-B (myh10) expression analysis during zebrafish embryonic development. Gene Expr Patterns 13, 265-270.

Hutchinson, S.A., Cheesman, S.E., Hale, L.A., Boone, J.Q., and Eisen, J.S. (2007). Nkx6 proteins specify one zebrafish primary motoneuron subtype by regulating late islet1 expression. Development 134, 1671-1677.

Jacob, J., Storm, R., Castro, D.S., Milton, C, Pla, R, Guillemot, R, Birchmeier, C, and Briscoe, J. (2009). Insm1 (IA-1) is an essential component of the regulatory network that specifies monoaminergic neuronal phenotypes in the vertebrate hindbrain. Development 136, 2477-2485.

Jia, S., Ivanov, A., Blasevic, D., Muller, T., Purfurst, B., Sun, W., Chen, W, Poy, M.N., Rajewsky, N., and Blrchmeler, C. (2015a). Insm1 cooperates with Neurod1 and Foxa2 to maintain mature pancreatic beta-cell function. EMBO J 34,1417-1433.

Jia, S., Wildner, H., and Birchmeier, C. (2015b). Insm1 controls the differentiation of pulmonary neuroendocrine cells by repressing Hes1. Dev Biol 408,90-98.

Lan, M.S., and Breslin, M.B. (2009). Structure, expression, and biological function of INSM1 transcription factor in neuroendocrine differentiation. FASEBJ 23, 2024-2033.

Landmesser, L.T. (1980). The generation of neuromuscular specificity. Annu Rev Neurosci 3, 279-302.

Levin, E.D., Aschner, M., Heberlein, U., Ruden, D., Welsh-Bohmer, K.A., Bartlett, S., Berger, K., Chen, L, Corl, A.B., Eddins, D., et al. (2009). Genetic aspects of behavioral neurotoxicology. Neurotoxicology 30, 741-753.

Lewis, K.E., and Eisen, J.S. (2003). From cells to circuits: development of the zebrafish spinal cord. Prog Neurobiol 69, 419-449.

Liu, D.W., and Westerfield, M. (1990). The formation of terminal fields in the absence of competitive interactions among primary motoneurons in the zebrafish. J Neurosci 10, 3947-3959.

Lorenzen, S.M., Duggan, A., Osipovich, A.B., Magnuson, M.A., and Garcia-Anoveros, J. (2015). Insm1 promotes neurogenic proliferation in delaminated otic progenitors. Mech Dev 138 Pt 3, 233-245.

Lukowski, C.M., Ritzel, R.G., and Waskiewicz, A.J. (2006). Expression of two Insm1-like genes in the developing zebrafish nervous system. Gene Expr Patterns 6, 711-718.

Mathelier, A., Fornes, O., Arenillas, D.J., Chen, C.Y., Denay, G., Lee, J., Shi, W., Shyr, C, Tan, G., Worsley-Hunt, R., et al. (2016). JASPAR 2016: a major expansion and update of the open-access database of transcription factor binding profiles. Nucleic acids research 44, D110-115.

Myers, P.Z. (1985). Spinal motoneurons of the larval zebrafish. J Comp Neurol 236, 555-561.

Myers, P.Z., Eisen, J.S., and Westerfield, M. (1986). Development and axonal outgrowth of identified motoneurons in the zebrafish. J Neurosci 6, 2278-2289.

Niederlander, C., and Lumsden, A. (1996). Late emigrating neural crest cells migrate specifically to the exit points of cranial branchiomotor nerves. Development 122, 2367-2374.

Osipovich, A.B., Long, Q., Manduchi, E., Gangula, R., Hipkens, S.B., Schneider, J., Okubo, T, Stoeckert, C.J., Jr., Takada, S., and Magnuson, M.A. (2014). Insm1 promotes endocrine cell differentiation by modulating the expression of a network of genes that includes Neurog3 and Ripply3. Development 141, 2939-2949.

Palaisa, K.A., and Granato, M. (2007). Analysis of zebrafish sidetracked mutants reveals a novel role for Plexin A3 in intraspinal motor axon guidance. Development 134, 3251-3257.

Park, H.C., Mehta, A., Richardson, J.S., and Appel, B. (2002). olig2 is required for zebrafish primary motor neuron and oligodendrocyte development. Dev Biol 248, 356-368.

Qi, J., Dong, Z., Shi, Y., Wang, X., Qin, Y., Wang, Y, and Liu, D. (2016). NgAgo-based fabplla gene knockdown causes eye developmental defects in zebrafish. Cell research 26,1349-1352.

Ramachandran, R., Zhao, X.F., and Goldman, D. (2012). Insm1a-mediated gene repression is essential for the formation and differentiation of Muller glia-derived progenitors in the injured retina. Nat Cell Biol 14,1013-1023.

Rodino-Klapac, L.R., and Beattie, C.E. (2004). Zebrafish topped is required for ventral motor axon guidance. Dev Biol 273, 308-320.

Shirasaki, R., and Pfaff, S.L. (2002). Transcriptional codes and the control of neuronal identity. Annu Rev Neurosa 25, 251-281.

Tanaka, H., Maeda, R., Shoji, W, Wada, H., Masai, I., Shiraki, T, Kobayashi, M., Nakayama, R., and Okamoto, H. (2007). Novel mutations affecting axon guidance In zebrafish and a role for plexin signalling in the guidance of trigeminal and facial nerve axons. Development 134, 3259-3269.

Wang, X., Ling, C.C., Li, L., Qin, Y., Qi, J., Liu, X., You, B., Shi, Y, Zhang, J., Jiang, Q., et al. (2016). MicroRNA-10a/10b represses a novel target gene mib1 to regulate angiogenesis. Cardiovascular research 110,140-150.

Westerfield, M., McMurray, J.V., and Eisen, J.S. (1986). Identified motoneurons and their innervation of axial muscles in the zebrafish. J Neurosa 6, 2267-2277.

Wildner, H., Gierl, M.S., Strehle, M., Pla, P, and Birchmeier, C. (2008). Insm1 (IA-1) Is a crucial component of the transcriptional network that controls differentiation of the sympatho-adrenal lineage. Development 135, 473-481.

Xie, J., Cal, T, Zhang, H., Lan, M.S., and Notkins, A.L. (2002). The zinc-finger transcription factor INSM1 is expressed during embryo development and interacts with the Cbl-associated protein. Genomics 80, 54-61.

Xu, M., Liu, D., Dong, Z., Wang, X., Wang, X., Liu, Y, Baas, P.W., and Liu, M. (2014). Kinesin-12 influences axonal growth during zebrafish neural development. Cytoskeleton (Hoboken) 71, 555-563.

Zelenchuk, T.A., and Bruses, J.L. (2011). In vivo labeling of zebrafish motor neurons using an mnx1 enhancer and Gal4/UAS. Genesis 49, 546-554.

